# Homeostatic Reinforcement Theory Accounts for Sodium Appetitive State- and Taste-Dependent Dopamine Responding

**DOI:** 10.1101/2023.01.26.525642

**Authors:** Alexia Duriez, Clemence Bergerot, Jackson J. Cone, Mitchell F. Roitman, Boris Gutkin

## Abstract

Seeking and consuming nutrients is essential to survival and maintenance of life. Dynamic and volatile environments require that animals learn complex behavioral strategies to obtain the necessary nutritive substances. While this has been classically viewed in terms of homeostatic regulation, where complex nutrient seeking behaviors are triggered by physiological need, recent theoretical work proposed that such strategies are a result of reinforcement learning processes. This theory also proposed that phasic dopamine (DA) signals play a key role in signaling potentially need-fulfilling outcomes. To examine potential links between homeostatic and reinforcement learning processes, we focus on sodium appetite as sodium depletion triggers state and taste dependent changes in behavior and DA signaling evoked by sodium-related stimuli. We find that both the behavior and the dynamics of DA signaling underlying sodium appetite can be accounted for by extending principles of homeostatic regulation into a reinforcement learning framework (HRRL). We first optimized HRRL-based agents to model sodium-seeking behavior measured in rats. Agents successfully reproduced the state and the taste dependence of behavioral responding for sodium as well as for lithium and potassium salts. We then show that these same agents can account for the regulation of DA signals evoked by sodium tastants in a taste and state dependent manner. Our models quantitatively describe how DA signals evoked by sodium decrease with satiety and increase with deprivation suggesting that phasic DA signals and sodium consumption are down regulated prior to animals reaching satiety. Lastly, our HRRL agents also account for the behavioral and neurophysiological observations that suggest mice cannot distinguish between sodium and lithium containing salts. Our HRRL agents exhibited an equal preference for sodium versus lithium containing solutions, and underestimated the nutritional value of sodium when lithium was concurrently available. We propose that animals use orosensory signals as predictors of the internal impact of the consumed good and our results pose clear targets for future experiments. In sum, this work suggests that appetite-dirven behavior may be driven by reinforcement learning mechanisms that are dynamically tuned by homeostatic need.

## Introduction

Seeking and consuming nutrients is essential to survival and maintenance of life. Animals living in dynamic and volatile environments must develop complex behavioral strategies to obtain the necessary nutritive substances. This has been classically viewed in terms of homeostatic regulation, where complex nutrient seeking behaviors are triggered by physiological need. Animals also seek nutrients in advance of acute need. How animals acquire nutrient-directed behaviors has most often been examined through the lens of reinforcement learning (RL) theories. In RL, subjects acquire information about signals from the environment that are associated with the receipt of reward (Rescorla & Wagner 1972). Importantly, RL signals are distributed throughout the brain (Tian et al 2016, Schultz 2000). Similarly, physiological need impacts a wide-array of brain circuits that regulate behaviors motivated by nutrient rewards (Livneh et al 2017, Geerling and Loewy 2008, Augustine, Lee, and Oka 2020). Intriguingly, the vast majority of RL theories do not treat the physiological origins of primary reward-seeking, nor do they speak to how nutrients and their associated values are modulated by internal state. To maximize survival, physiological needs should augment signals that drive RL to promote learning in environments that offer access to essential nutrients. Thus, the reinforcing value of a nutrient, and consequently the degree to which an RL-based agent can learn from actions that acquire said nutrient, should be modulated in an appetite-dependent manner.

An essential area for exploration is thus the degree to which homeostatic and reinforcement learning processes are coupled in the central nervous system. RL processes have been most closely associated with mesolimbic circuitry, namely the midbrain DA neurons and their major target, the striatum (Schultz 1998, Keiflin and Janak 2015, Dabney et al. 2020). While debate remains as to the role of DA in RL (Kutlu et al. 2021), it is increasingly clear that midbrain DA neurons and their responses to essential nutrients are modulated by physiological state, through direct hormonal influence (Liu and Borgland 2015, Cone et al. 2015, Mietlicki-Baase et al. 2015, Mebel et al. 2012) or via interactions with homeostatic and/or related circuits (Cone et al. 2014; Cone et al. 2016, Fortin and Roitman 2018; Hsu et al. 2020; Grove et al. 2022; Reichenbach et al. 2022). A particularly powerful example of the impact of physiological need on motivated behavior and DA signaling is sodium appetite. Sodium appetite is a natural behavior (Denton 1982) whereby a sodium deficit generates sodium-seeking behaviors and selective consumption of sodium over other nutrients. Under homeostatic conditions, rodents avoid consumption of hypertonic sodium solutions. However, sodium depletion (via injection of a diuretic/natriuretic, e.g., furosemide) or removal of the adrenal glands (Richter 1936) induces avid consumption of hypertonic sodium solutions and appetitive taste reactivity (Berridge et al. 1984). Importantly, phasic DA responses to the taste of a hypertonic sodium solution are dynamically sensitive to sodium balance (Cone et al. 2016, Fortin and Roitman 2018). As with behavior, the DA response in sodium deplete rats is blocked by lingual application of the epithelial sodium channel blocker amiloride (Cone et al. 2016) and is selective for sodium solutions (Fortin and Roitman 2018). Lithium chloride, a notable exception, is equally preferred (Nachman, 1963; Fortin and Roitman 2018) likely due to sodium taste fibers responding to lithium as well (but not potassium) (Contreras et al. 1984). These data argue strongly that information related to the current state of sodium balance is communicated to midbrain DA neurons to regulate brain signals thought to drive RL. Taken together, these data pose a major challenge to current state of the art RL theories and novel RL models need to be developed that account for the impact of physiological need and the role of gustatory information on reward learning. Sodium appetite is an ideal paradigm to address this issue.

We recently put forth a Homeostatic Reinforcement Learning (HRRL) framework that was developed to study how animals learn need-based adaptive behavioral strategies in their environment to obtain rewarding outcomes (Keramati and Gutkin, 2014). The HRRL agent learns to maximize the total cumulative reward by performing actions and predicting the impact of their outcome on its internal state. This framework relies on a new definition of rewards: the rewarding value of an action is a function of the predicted impact on the difference between the current internal state and the ideal one (i.e., “setpoint”). The function that links the internal states to rewards is called the drive-function. In other words, the reinforcing value of a stimulus is modulated by the degree to which it alleviates or exacerbates a physiological need. In this way, HRRL joins the predictive homeostatic regulation and reinforcement learning theories by positing that minimizing deviations from a homeostatic set point and maximizing reward are equivalent. In other words HRRL synthesizes RL algorithms with the drive-reduction theories of motivation (Hull 1943). HRRL has been used to simulate the consumption of various resources and reproduce experimental data. It can also be used to represent complex behavior such as anticipatory responding, binge-eating (Keramati and Gutkin, 2014) and cocaine addiction (Keramati et al., 2017). Interestingly, it can be shown mathematically that HRRL agents show predictive allostatic behavior and HRRL accounts for the incentive salience proposals: the internal state of the HRRL agents is dynamically changed according to upcoming challenges and the action values (incentives) are modulated dynamically by the internal state of the animal.

Here, we show that the HRRL model can account for sodium-seeking behavior and DA signaling in rats. We first required the HRRL models to reproduce behavioral data showing that sodium-deprived rats preferred sodium and lithium over potassium solutions. We then showed that such HRRL agents also reproduce the dynamics of DA signals. We then used the models to make several predictions about satiety dependent modulation of behavior and how exposure to lithium only may modulate the behavior and the reinforcing value sodium.

## Methods

### HRRL theory for sodium consumption

#### State space representation

The internal state is considered as a continuous variable that can be represented at each time t by a point in a homeostatic state space. As theorized by Keramati and Gutkin (2014), this state space has one dimension per homeostatic variable. The ideal internal state is the equilibrium point of the homeostatic state space. This point we call the setpoint represents the internal state that maximizes the chances of survival (satiety). It is denoted by H*= (h1 *, h2 *,…, hN *). In this study, the state space has only one dimension, corresponding to the internal sodium level.

#### Reward calculation mechanism

The HRRL theory provides a function called the drive, which takes as its argument the degree of departure from a “satiety” point and has a unique minimum at that set point. The drive is a function of the deviation of the animal’s internal state from its homeostatic setpoint *H\** (Figure 1). In a homeostatic state space with one dimension, the drive is given by the following expression (Keramati & Gutkin, 2014):

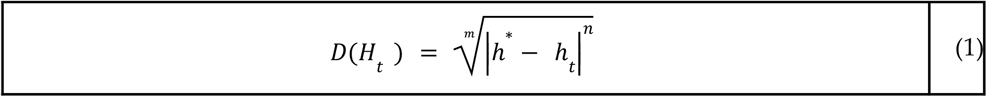

where *t* represents the time, *m* and *n* are free parameters that influence non-linearly the mapping between homeostatic deviations and the rewarding value of their reduction.

**Figure 1.**
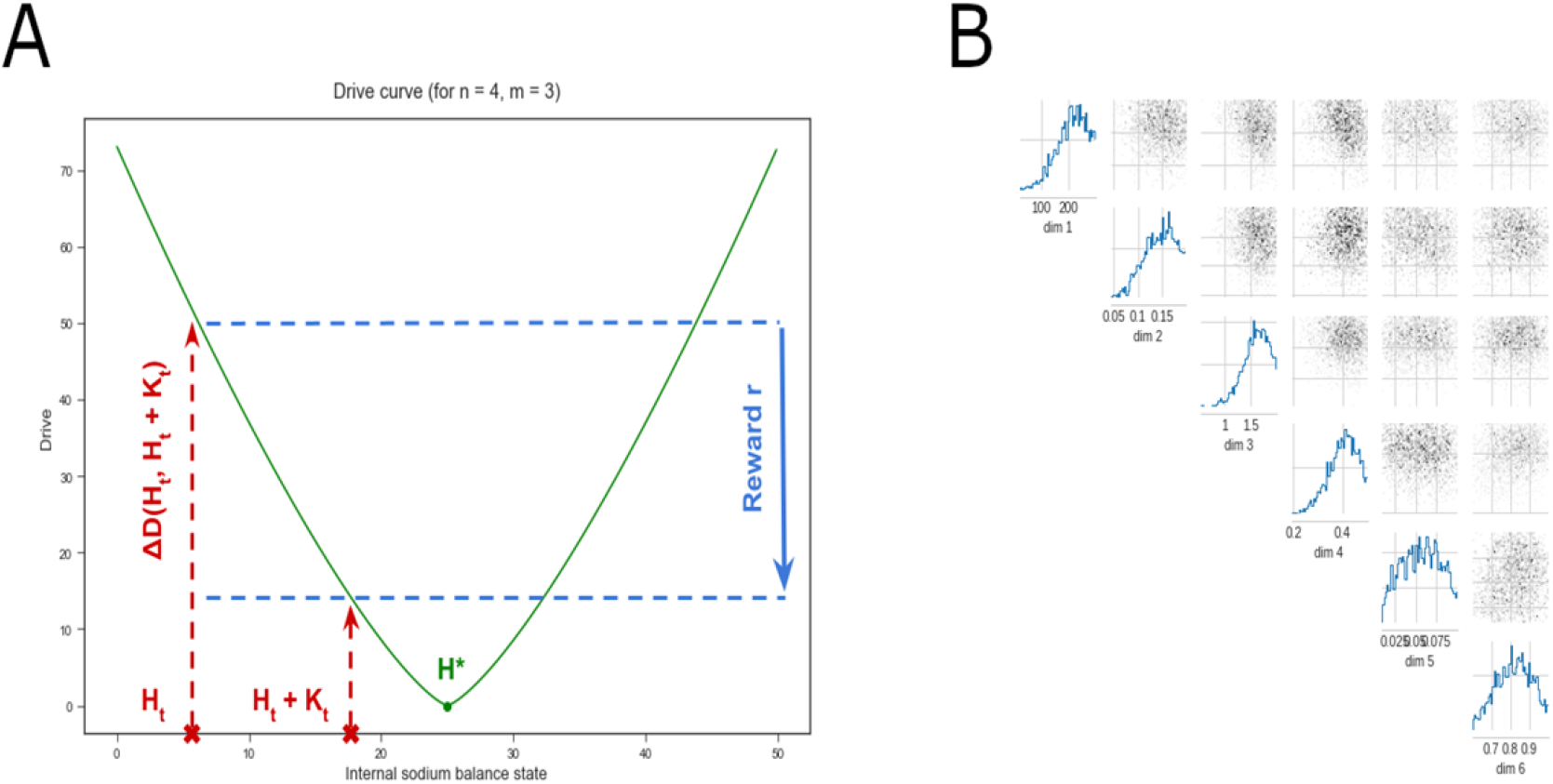
A: Drive as a function of the internal sodium level. The optimal sodium level is denoted by H*, indicated by a dot. The current sodium state of the agent at time t is Ht. An action apports an outcome that impacts the sodium level denoted by *k_t’_*, transitioning the internal state to Ht+1=Ht+Kt. The change in the drive function is defined as the reward r=ΔD(Ht, Ht+Kt). B: Probability distributions for parameter values yielding model results consistent with the experimental observations. Parameters are, from top to bottom: the setpoint *H\**, the learning rate *ϵ*, the rate of exploration *β*, the fixed outcome K, the loss of sodium after each trial and the energy cost of drinking. If each parameter takes a value with a high probability, the simulation results are consistent with the experimental observation. For each pair of parameters, the joint probability distribution that the two parameters fall in their respective range of possible values is also represented. Distributions obtained following Gonçalvez et al (2020), see Methods for details.

As an animal performs an action, its internal state is modified by the outcome *K_t_* of the action. The homeostatic reward is defined in a non-circular way, as the reduction of the homeostatic distance from the setpoint caused by the outcome *K_t_*.

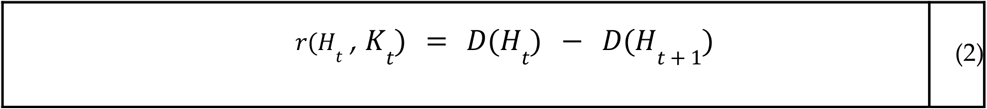

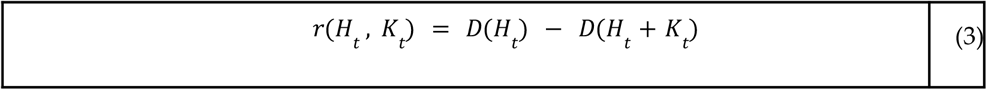

The reward associated with taking an action from state *H_t_* resulting in an outcome *K_t_*. that transitions the internal state to *H*_*t*+1_ is positive if the subsequent internal state (*H*_*t*+1_) remains below or equal to the setpoint However, if the animal is currently at its setpoint (i.e., *H_t_* = *H*\*), the reward value obtained with the outcome *K_t_*. is negative: the outcome is negatively reinforcing.

#### Taste value estimation mechanism

We hypothesize that animals sense the rewarding value of a tastant through the gustatory information they receive before experiencing its postingestive qualities (Keramati and Gutkin, 2014). The reward is thus computed by the animals orosensory approximation of the nutritional value *K_t_* given the amount of solution they consumed. This estimate of the outcome, based on the orosensory properties of the stimulus, is denoted by 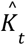. The reward therefore becomes:

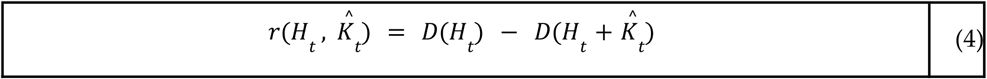

We further hypothesize that 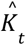 is not constant. In our model, 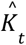 is learned with a learning rate ϵ through tasting the corresponding solution and experiencing its impact on the internal sodium level *H_t_*. We introduced this aspect since it has been suggested that assigning a reinforcing value to a taste of food requires that the animal experiences its nutritional impact (Beeler et al., 2012). According to this study, hungry animals learn that a taste stimulus is predictive of a need-reducing reward by experiencing the association between these two properties of food. We consider that 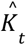 also has an innate non-zero initial value, which is supported by recording of DA transients: the first NaCl infusion already elicits a DA response (Cone et al. 2016).

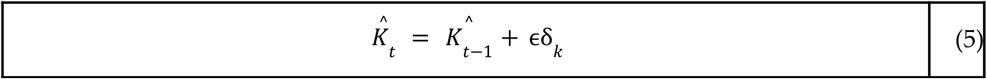

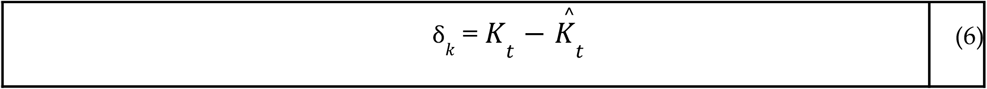

It has been shown that in the absence of prior experience, sodium-depleted rats cannot discriminate between sodium and lithium chloride (Fortin and Roitman 2018). We therefore hypothesize that taste information alone is insufficient to distinguish between sodium and lithium in the absence of any post-ingestive impact on *D*. In our model, this means that, for naïve animals, sodium and lithium have the same 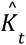, which represents the estimated impact of a sodium-like tasting solution on the internal state.

#### Action value estimation mechanism

With the Q-learning method, rats estimate the value *v* of each choice as they discover which actions are more rewarding than the others (Sutton & Barto, 1998). Once a rat executes an action *a* and the homeostatic reward *r* is computed, the value *v*(*a*) of this action is updated using the reward prediction error (RPE), δ*_r_*, with the learning rate ϵ (Keramati and Gutkin, 2014).

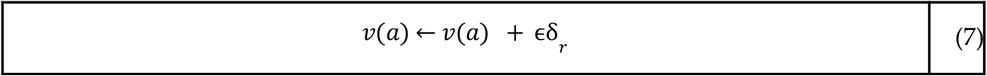

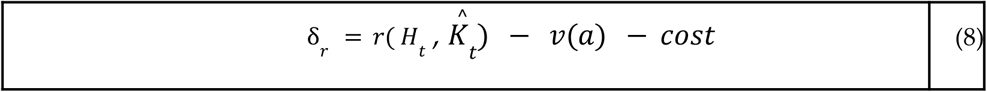

The cost is a penalty we introduced in this model associated with the energy cost of approaching the sipper tubes and consuming any of the solutions. By decreasing the reward prediction error term (RPE), it reduces the reinforcing value of an action, and thus the motivation of the agent to pursue that action in the future. We assume that the cost of approaching and drinking from a sipper tube is apriori encoded in the rats’ representation of their environment.

δ*_r_* is the RPE signal which is purportedly encoded by midbrain dopaminergic neurons (e.g. see Schultz et al., 1997). We therefore monitor the RPE by sampling DA fluctuations in the Nucleus Accumbens (NAc). A positive RPE in our model corresponds to a phasic DA response in the NAc. The RPE signal is negative when the predicted reward is superior to the actual one. The RPE can also be negative for actions that yield no reward due to the energy cost associated with performing said action.

#### Action choice mechanism

The probability of taking an action *a_i_* is proportional to its estimated value *v*(*a_i_*), according to the softmax rule (Sutton & Barton, 1998):

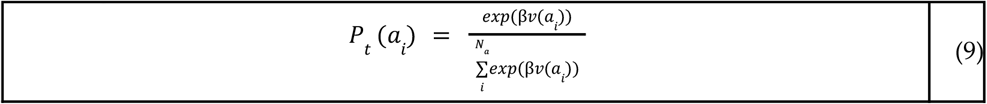

*N_a_* is the number of possible actions, such as drinking sodium chloride or drinking nothing. The probability distribution over the possible actions is the stochastic policy used by the simulated rats to choose their next action. *β* is a parameter used to modulate the probability of selecting actions with different estimated values. A high beta causes the selection probabilities to diverge faster. For extreme betas, this can make the action choice almost deterministic (i.e., greedy). On the contrary, a low beta makes the probabilities change more slowly, and actions are therefore selected more randomly.In general *β* controls exploration versus exploitation behaviors.

#### Trial schedule

A new trial begins every 15 seconds. At the beginning of a trial, the rats choose an action based on the learned stochastic policy. The outcome of the action is added to the internal state.

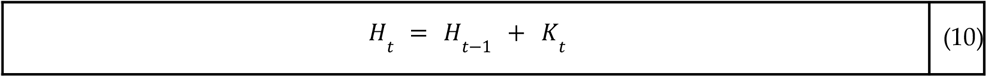

The new drive, then the homeostatic reward, is calculated. The RPE is computed, then the value of the action performed is updated. Meanwhile, the values of the other actions remain unchanged. The error δ*_k_* in the approximation of *K_t_* is also computed, allowing 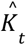 to be updated. The policy is then updated using the new action values. Finally, the internal state loses a small quantity of sodium, to simulate grossly the dynamics of sodium in a real organism:

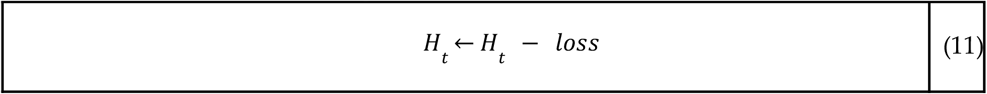

#### Optimal parameters

We first used an ad hoc procedure to adjust the free parameters of the model to yield a qualitative agreement with the data and to determine physiological ranges that would indeed produce such appropriate model behavior. The “Simulation-Based Inference” approach was then employed to find optimal parameter distributions, within a range of possible biologically values estimated with the search by hand, for the algorithm to return a desired output (Gonçalvez et al, 2020 [14]). The parameters used in the simulations in the manuscript were then sampled from the modes of those distributions to produce individual simulated animals (agents).

For clarity we briefly review the SBI methodology. The Simulation-Based Inference method uses an algorithm called “Sequential Neural Posterior Estimation” (SNPE). SNPE requires three inputs (Gonçalvez et al. 2020):

- A simulator, which is the learning algorithm (an HRRL agent), returning one output of interest. The preference score of potassium chloride when sodium and potassium are available was chosen as the output of the simulator.
- Prior knowledge on the free parameters, in the form of a uniform distribution over the possible values each can take, within a defined interval. The maximum and minimum values each parameter can take are chosen based on biological plausibility, previous models and the set of values found by hand.
- The observations of the experimental data, which we want to reproduce as closely as possible with our model, is the preference score for potassium relative to sodium at the end of a 10-minute two-bottle intake test that should match the data to be 0.1.

SNPE returns for each parameter a probability distribution over its range of possible values. The values with a high probability are consistent with the observation. Simulation-Based Inference was used to determine the values of the setpoint H*, the learning rate ε, the rate of exploration *β*, the fixed outcome K of every drink taken by the rats, the loss and the cost. 2000 samples were drawn from 1000 batches of 1000 simulations via the SNPE method to obtain the distributions Figure 1B. Each parameter was assigned the average value of the distribution, as an approximation for the value with the highest probability.

For each simulated agent, we recorded the evolution of multiple parameters or values: 1- the probabilities of consuming one solution over another 2-the amount of ingested sodium chloride 3- the predicted nutritive value associated with the taste of sodium, 4- the reward prediction error signal, and 5-the choices made throughout the simulated experiments. Individual agents were endowed with parameters that were sampled from the peaks of the posterior distributions (see above), hence representative instantiations of optimal parameter sets. The agents are then used to show how the studied variables evolve depending on the solutions available and the initial internal state. Interagent variability was introduced to calculate the statistics (e.g. averages) of the relevant variables over several different individual agents (to match the experimental data). To generate individual agents, we sampled the free parameter distributions within the ranges giving the highest probability of the agreement with the data. This way we obtained a simulated cohort of animals - model agents, for which we could compile response statistics. The free parameters of the HRRL model are the setpoint, the learning rate, the exploration rate and the loss of sodium between each trial. For each of these parameters and individual agent, values were randomly sampled within a range in which the probability of the learning algorithm output to be consistent with the behavioral data from Fortin and Roitman (2018) is around its maximum (approx. 1 std of the peak, see Figure 1B).

### Statistical analysis

In the simulated experiments, several variables were calculated in 14 simulated rats having access to potassium and 15 simulated rats having access to lithium: the probability of choosing each of the available solutions at the end of the experiment, the average reward prediction error throughout the experiment, the predicted nutritive value of one lick of NaCl or LiCl (*K_t_*) and the number of trials necessary to reach the optimal sodium level. For each of these variables, the averages of the two groups were compared with a two-tailed Welch’s t-test. The preference scores for potassium group (n=14) and lithium group (n=15) were compared using a two-tailed Student’s t-test. The cumulative consumption between the two groups were compared with a two-tailed Student’s t-test. This analysis aims to reproduce analysis methods used by Fortin and Roitman’s methods (2018).

## Results

We first optimized the free parameters of the model (see Methods) to generate a cohort of individual agents whose simulated behavior reproduces the results of the 10-minute two bottle intake test conducted by Fortin and Roitman (2018). The parameters sampled and optimized were the setpoint, the learning rate, the exploration rate and the quantity of sodium lost between each trial. They were tuned for the potassium preference score of the simulated sodium-deprived agents to be as close as possible to the experimentally observed values. Our goal was to capture the preference scores for the different salt solutions as a function of the animal’s internal state and choices in the experiment. To illustrate the results, we picked a parameter set (see Annex 1 Table 1) with nearly optimal parameter values producing a simulated individual “rat” (N.B. for the rest of the manuscript we will denote such simulated animal by agent). **Figure 2** shows the behavior of such an individual agent throughout the experiments, and indicates that the model accounts for the qualitative observations made by Fortin and Roitman (2018) on the preference for sodium and lithium over potassium. In **Fig2A** the evolution of the probability for an agent with optimized free parameters to consume either NaCl, KCl or nothing (left column), and the evolution of the probability for the same agent to consume either NaCl, LiCl or nothing (right column). In both simulations the agent was initialized with a simulated depleted sodium internal state variable. As expected, when sodium and potassium are available, the probability of consuming NaCl increases greatly while the probability for KCl consumption decreases and becomes lower than the chances of drinking nothing. This suggests that sodium-deplete agents selectively target the taste of sodium in their strategies to regulate their sodium level (as seen in the experiment). On the other hand, the probability of consuming NaCl or LiCl remains around 0.4, higher than the probability of drinking nothing. This indicates that the taste of sodium and lithium are both expected to restore sodium balance.

**Figure 2.**
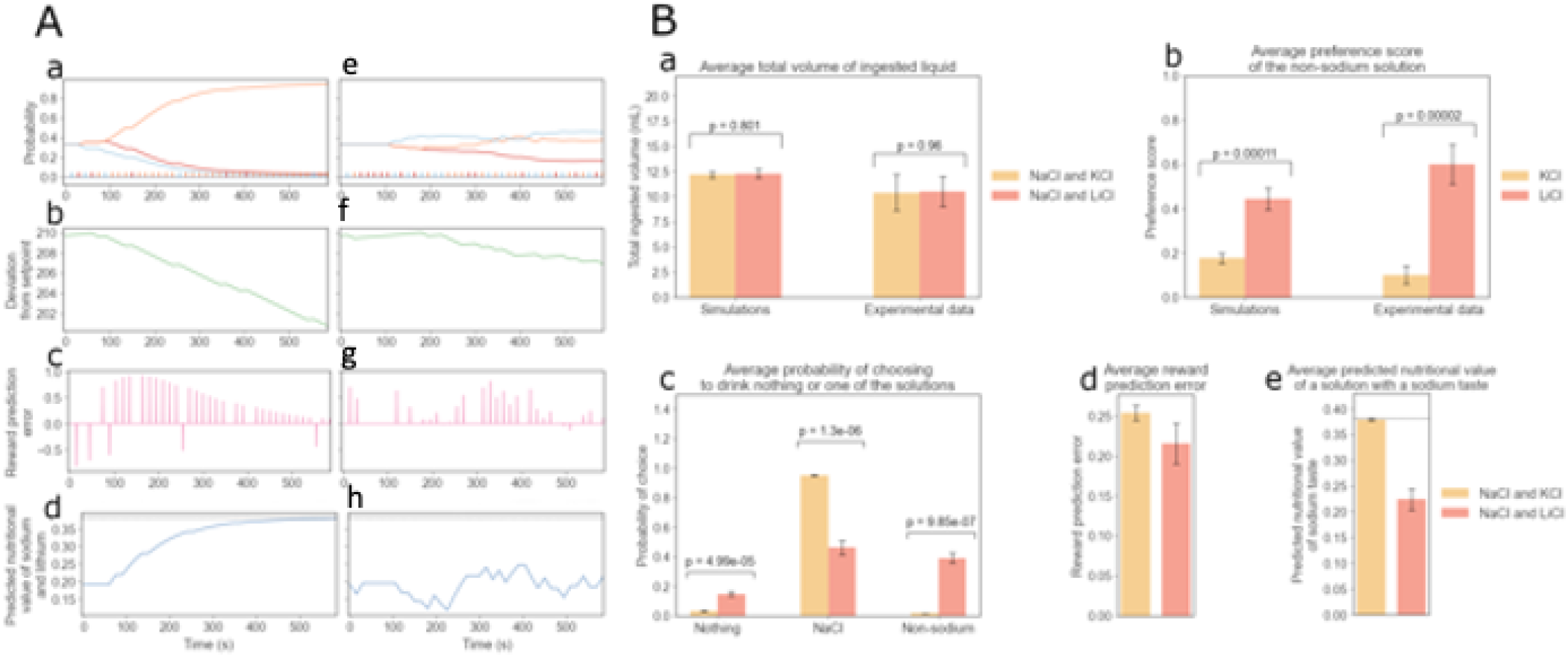
Simulations for the two-bottle intake test (Fortin and Roitman, 2018) are consistent with experimental observations on sodium appetite. Simulation results are shown for experiments where sodium-depleted rats have access to NaCl and KCl (left column, Aa-d), and in the other version, NaCl and LiCl are the solutions available (right column, Ae-h). Both versions of the experiment last 10 minutes (40 trials). **A (left column):** Aa: Evolution of the choice probability for a simulated agent with optimized free parameters to consume NaCl, KCl or nothing over time (left). Evolution of the probability for an agent with optimized parameters to consume NaCl, LiCl or nothing over time (right). Note that the two simulated individuals are identical in the values of the free parameters: the discrepancy between their behaviors is due to the different solutions available to them. Note that the bars at the bottom indicate the individual choices (color codes the choice). A**b,** Amount of sodium ingested over time. Note that for these simulations, the optimal level of internal sodium for survival is set at 212 in arbitrary units. **Ac:** Reward prediction error signals of over time. Note that in our simulations the RPE is a proxy for the phasic dopamine signal. **Ad:** Evolution of the approximated nutritional value 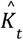, for sodium and lithium. **Ae-h**: panels show results corresponding to the left column but with NaCl and LiCl as choices.

**Table 1.** Free parameters of the model and their values.

| Parameter | Value | Explanation |
| --- | --- | --- |
| $\epsilon$ | 0.1446 | Learning rate |
| $\beta$ | 1.5896 | Rate of exploration |
| m | 3 | Root of the Drive function |
| n | 4 | Power of the Drive function |
| K | 0.3819 | Outcome of an action and its effect on the internal sodium level |
| cost | 0.8180 | Energy cost of going to drink from a sipper tube |
| loss | 0.0559 | Quantity of sodium metabolized by rats at every trial |
| setpoint $H^*$ | 211.7066 | Optimal sodium internal level |

The individual choices made by the agent are also shown in **Figure2 Aa and Ae**. This allows us to understand how the preference for sodium relative to a non-sodium salt evolves depending on the learned subjective value of each solution and the deviation from the setpoint (**Figure2 Ab and Af**). We can make an inference about the dynamics of DA signaling in the NAc by studying the reward prediction errors (RPEs) of the agents during the simulated experiments (**Figure2 Ac and Ag**). Notice that NaCl and LiCl consumption gives rise to positive RPE signals, which correspond in our model to the experimentally observed positive DA responses in the NAc. Note that in the NaCl/LiCl task, the RPE signals are rapidly quenched, as the agent rapidly learns that both of these outcomes have an equal orosensory quality. We also tracked how the approximation of sodium nutritive value may evolve throughout the experiments (**Figure2 Ad and Ah**). We can see that when NaCl and KCl are present the value of NaCl choice increases (the value of KCl decreases, result not shown), whereas in the NaCl/LiCl task the value of NaCl stays relatively constant since their orosensory quality is equal. Indeed, our HRRL agents simulations show how DA dynamics are linked to internal state fluctuations, and also how the animals’ learned estimation of the nutritional value is based on gustatory information.

The model reproduced quantitatively the experimental results (Fortin and Roitman 2018) for the amount of ingested liquid and the individual preference for sodium, lithium or potassium. **Figure 2**: **Ba** and **Bb** show that the average amount of liquid ingested during the experiment and the preference scores for the non-sodium solutions obtained with the simulations are consistent with the experimental values. We further confirmed qualitative observations for simulated individual agents (shown in **Figure 2A**): at the end of the 10-minute experiment, rats (as well as the simulated agents) with access to NaCl and LiCl (N = 15) are approximately equally likely to drink either of the two solutions. Rats with access to NaCl and KCl (N = 14), however, are unlikely to drink KCl, with a choice probability close to zero (see **Figure 2 Bc**).

Based on the behaviorally validated model, we asked what would be predicted for the dopaminergic signal (in our model this corresponds to the PRE) and the predicted value associated with the sodium gustatory cue during the final part of the experiment. **Figure 2Bd** shows that the inter-individual-average phasic DA signal strength (as measured experimentally by the DA concentration) in the NAc during the 10-minute experiment is higher when KCl is available than when LiCl is present as a choice, while in **Figure 2Be** we show the average predicted value of a lick of sodium (or lithium) associated with the corresponding taste for the model agents at the end of the 10-minute simulated experiment. The true nutritive value of one lick of sodium is indicated with a dotted line. Our simulations predict that rats learn the true value of sodium associated with its taste when NaCl and KCl are available. However, their estimate is equal to about half of the true value when NaCl and LiCl are available. This higher value of the rats’ estimated value of sodium and lithium when KCl is available may explain why the average RPE is also higher in this condition as compared to the LiCl-available condition (**Panel Bd**). On the other hand, when NaCl and LiCl are both available, the taste-dependent rewards are equally distributed between the two solutions and hence the RPE for NaCl is reduced and a positive RPE is misattributed to the LiCl choice (in the sense that it is based purely on the taste information and not on the internal impact). Hence LiCl acquires a positive predicted value at the expense of the NaCl choice.

We then set out to see if the behavior-optimized model could also account for experimentally observed NAc DA responses to intra-oral NaCl delivery in sodium-depleted rats and how this response develops as the animal continues to consume NaCl. Figure 3 shows a simulated agent with parameters optimized to account for the experimental behavioral preference scores, as in the previous simulations. We can see that the simulated agents’ RPE signals are qualitatively consistent with the experimentally observed DA responses. Cone et al. (2016) measured the average DA response evoked by ten intra-oral infusions of NaCl in sodium-deplete rats (N = 5). They observed that the first five infusions evoke larger DA responses than the last five. Fig 3**A** shows the simulation results for the same experiment alongside the summary data from Cone et al. In order to simulate this experiment, we first measured the RPE for the naïve depleted model during the first 5 simulated trials of exposure to NaCl stimulus, then during the last 5 trials. The model was “depleted” by initializing the internal state significantly below the optimal value (variable *H_t_* = 2). NaCl delivery was simulated as a series of 10 trials where the agent receives the palatable sodium reward. In the model, the impact of NaCl on the internal state was simulated by a dose-dependent shift in the internal state variable (here of *K_t_* = 0.3819 per injection) and increase of the taste signal by ϵδ*_k_*, with 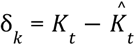 (see methods). The results were averaged over 5 agents whose free parameter values were randomly sampled as previously presented. To compare the RPE with biological results, we referred to Cone et al.’s experiments (2016). With their data, we calculated the baseline DA concentration for each trial and each rat as the average DA concentration during the 4 seconds preceding the NaCl infusion period. We subtracted this baseline from the DA concentration values during the 4-second infusion period. Then, for each rat, we computed the average peak of the DA concentration increase over the first and the last 5 trials. These two variables were finally averaged over the 5 rats. In order to compare the DA signal strength to the RPE, we normalized the DA signal from the first 5 trials and the RPE from the first 5 trials to unity. Consistent with the experimental results, we observed a decrease in the response between the first group of trials and the last. Indeed, Welch’s t-test showed that the difference between the average responses to the first 5 trials and to the last ones was significant (p = 6.93e-07). Moreover, the differences between the simulation and the experimental results were not statistically significant (Welch’s t-test; see p-values on the figure). We then asked if the model can account also for differential taste-dependence of the phasic DA response. Fortin and Roitman (2018) tracked the average DA response of sodium-deplete rats during 10 intraoral infusions of NaCl, of KCl, of LiCl or of water and found that NaCl and LiCl evoke a strong DA response while KCl does not. We simulated an analogue experiment. **Fig 3B** compares the simulated and experimental results. Note that in our simulations we left out the response to water since that would, in the simulations, lead to a null result. With the experimental data from Fortin and Roitman (2018), we computed the peak of the DA concentration during the 4-second infusion period and subtracted the average DA concentration measured during the 4 seconds before the infusion onset as a baseline. This baseline was computed for each rat. We then averaged the DA increase over the rats. For the simulated data, we computed the average RPE signal evoked by 10 infusions of each solution. The results were averaged over 6 or 5 agents whose free parameter values were randomly sampled as previously. The baseline is null in our model. Then in order to compare the DA data to the RPE we normalized the response to NaCl to unity. We show that in our simulations NaCl and LiCl are also the only solutions triggering a DA signal. Welch’s t-test showed that the differences between the simulation and experimental results are not statistically significant.

**Figure 3.**
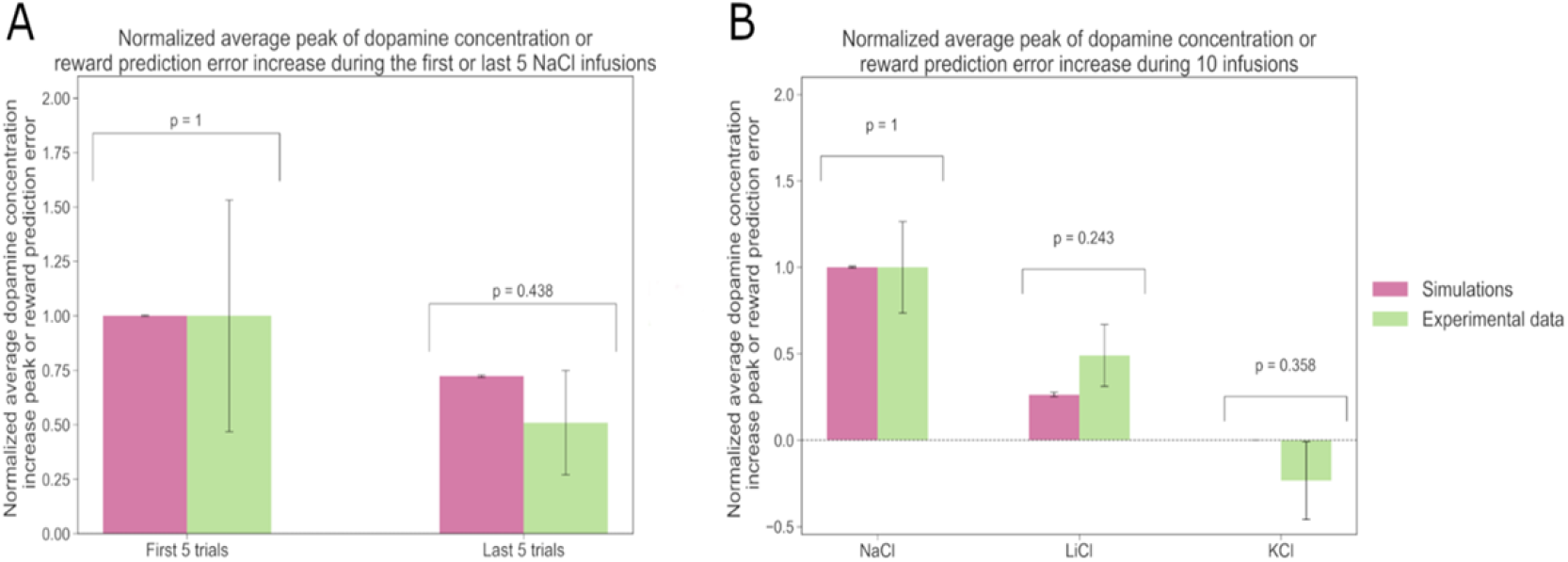
The model accounts for experimental data on dopamine dynamics. **A**, Figure taken reproduced from Cone et al.’s publication (2016): average dopamine response evoked by ten intra-oral infusions of NaCl in sodium-deplete rats (N=5). The average dopamine response to the first five infusions is shown in black and the response to the last five are in gray. **B**, Comparison of the sum of the dopamine concentration increase during the infusion period, averaged over 6 or 5 rats, and averaged over the first 5 or last 5 of a series of 10 intra-oral NaCl infusions, as reported by Cone et al. in 2016, (in green) with the average RPE during the first 5 or last 5 of 10 successive simulated infusion trials, averaged over 6 or 5 rats (in pink).

We further reasoned that original 10-minute two-bottle intake test used by Fortin and Roitman (2018) may not give enough time for the acutely sodium-deplete rats to reach sodium satiety (or in terms of our model – to reach the optimal settling point of its internal state that corresponds to the minimum of the drive function). In order to study rat behavior when their sodium internal state approaches and reaches the optimal balance, we simulated the two-bottle intake test over a period that was sufficiently long for the agent to reach the statiation state. **Figure 4A** shows simulations for the evolution of behavior and DA dynamics when the agents approach satiety and become replete in sodium. **Figure 4Aa,e** show the evolution of the probability for a simulated individual with optimized free parameters to drink either NaCl, KCl or nothing (a), and the evolution of the probability to drink either NaCl, LiCl or nothing (e). Interestingly, in both simulated conditions, reduction in consumption of NaCL or LiCL appears to anticipate satiety: the probability to drink starts to decrease before the amount of NaCl ingested (shown in **Figure 4Ab,f**) makes the individual reach satiety. We then tracked the RPE particularly when agents approach the replete state. In this case, consuming NaCl deviates the internal state away (beyond) from the setpoint. This deviation translates into a negative RPE. In our model, assuming that the baseline of DA outflow is sufficiently low, this corresponds to no DA being released in the NAc (should the baseline be high, the negative RPE would be interpreted as a phasic decrease of DA outflow. This is consistent with the fact that in sodium replete animals, intraoral infusions of hypertonic sodium chloride evoke no DA signal in the NAc: the DA concentration does not change from baseline (Fortin & Roitman, 2018 and Cone et al., 2016). We also note the tell-tale oscillatory pattern in the RPE/DA over trials for the replete condition. This may be explained as follows: a small amount of sodium is lost between each trial, therefore after having reached a replete state for the first time, as the agents have learned not to consume NaCl, the internal state can move below the setpoint again. When this happens, NaCl becomes rewarding again, which corresponds to the positive peaks of the RPE. The dynamics of RPE and the sodium-consumptive choice frequency is then determined by the physiological processes that regulate sodium loss.

**Figure 4.**
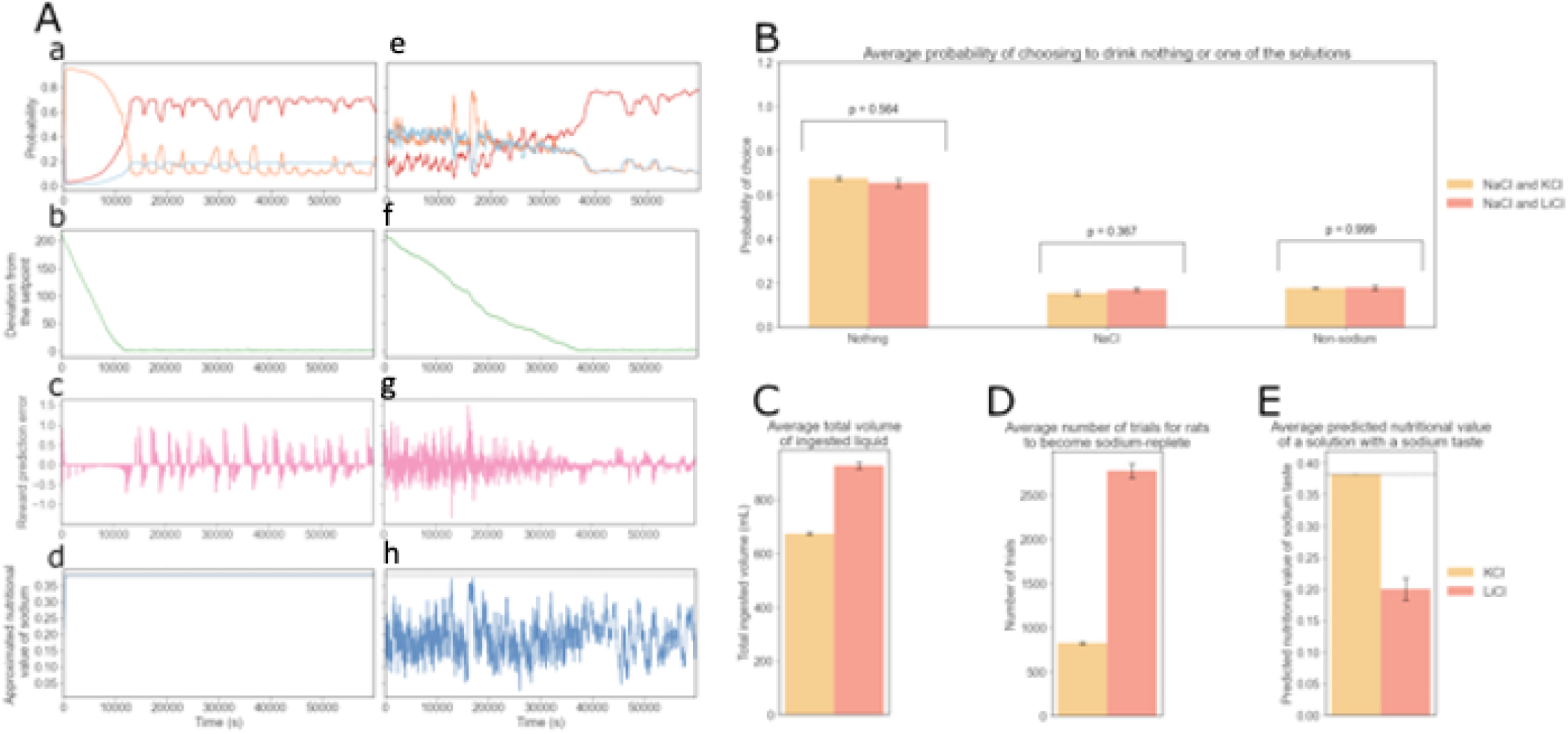
Model predictions for behavior during approach to sodium satiety. Here we produced example simulated data for two rats taking the two-bottle intake test developed by Fortin and Roitman (2018) for an extended period of time. In one version of the experiment, sodium-deplete agents have access to NaCl and KCl, and in the other version, NaCl and LiCl are the solutions available. Both versions of the experiment last 1000 minutes (4000 trials). For panels in B: 14 agents perform the version with KCl and 15 others perform the version with LiCl. The free parameter values for the agents in this group (N = 28) are all different and are sampled around the optimized ones used for the example data shown in A. Aa and e, Evolution of the probability for an optimized agent to drink NaCl, KCl or nothing over time (left). Evolution of the probability for the same agent to drink NaCl, LiCl or nothing over time (right). Ab and f: Deviation from the optimal sodium state. The optimal level of internal sodium for survival is set at 0. Ac and g: Reward prediction error signals. Ad and h: Evolution of the approximated nutritional value (K_t) ^ for sodium (d) and lithium (g). B, Average probabilities for agents (N = 28) to drink NaCl, the available non-sodium solution (KCl or LiCl), or nothing at the end of the 1000-minute experiment. Probabilities for agents that have access to NaCl and KCl (N = 14) are shown in yellow, and probabilities for access to NaCl and LiCl (N = 15) are shown in red. C, Average volumes of ingested liquid by agents (N = 28) during the 1000-minute simulated experiments. The intake volume for agents that have access to NaCl and KCl (N = 14) is shown in yellow, and the intake volume for access to NaCl and LiCl (N = 15) is shown in red (p = 1.39e-11). D, Average number of trials to reach the setpoint for sodium for the first time during the simulated experiments. The number of trials for agents that have access to NaCl and KCl (N = 14) is shown in yellow, and the number of trials for access to NaCl and LiCl (N = 15) is shown in red (p = 1.27e-11). E, Average predicted nutritional value of sodium or lithium associated to their taste at the end of the 1000-minute simulated experiments. This value for access to NaCl and KCl (N = 14) is shown in yellow, and for rats with access to NaCl and LiCl (N = 15) it is shown in red (p = 4.45e-07).

When there is access to lithium chloride in the replete state, consuming this solution does not change the sodium internal state, but it can still be a punishment, similar to drinking sodium, since the two solutions acquire the same predictive value due to taste similarity. As in the 10-minute experiment, **Panel 4Ad,h** indicates that the value of the predicted nutritional value of sodium associated to its taste, 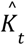 is learned when rats have access to NaCl and KCl. On the other hand, when rats consume both NaCl and LiCl, the long-term value of *k_t_* oscillates around the value learned in the first 10 minutes of the experiment (equal to half the real nutritional value of one lick of NaCl).

**Figure 4, right panels** show average results for this simulated experiment, that could be tested with new experiments. First, **Panel 4B** confirms the qualitative observation from Figure 4A that in the replete state, the probability to drink nothing is much higher than the probability to drink one of the two solutions (NaCl or LiCl). Interestingly, agents with access to NaCl and LiCl (N = 15) appear to be slightly more likely to consume NaCl than agents in the other condition (N = 14). This is likely because the agent’s estimation of the nutritive value of sodium (under LiCl-NaCl access) is lower than under NaCl-KCl access (**Panel 4E**). So consuming NaCl in a replete state under LiCl-NaCl access is a weaker punishment than for agents with access to NaCl and KCl. We are also interested in the amount of liquid agents consume during the experiment, in each condition (**Panel 4C**). It appears that more liquid is consumed when NaCl and LiCl are the solutions available. This can be linked to the fact that more trials are necessary for the agent to reach satiety in this condition (**Panel 4D**). Finally, in **Panel 4E**, the agents appear to learn the true value of sodium associated to its taste when NaCl and KCl are available. However, their estimate is equal to approximately half of the true value (indicated by a dotted line) when NaCl and LiCl are available.

Evidence suggests that gastric distension is an early inhibitory signal of water ingestion in thirsty rats (Hoffmann et al., 2006). Hoffmann et al show that dehydrated rats will almost continuously drink water or saline, yet stop drinking after 5 to 8 minutes before reaching satiety. The oropharynx also is likely to be involved in the anticipatory control of drinking behavior, by signaling the amount of water consumed (Zimmerman et al., 2017). Considering these results on thirst, we can hypothesize that animals would stop drinking before reaching sodium homeostasis because of early inhibitory signals. Arguably, an experiment where rats would consume sodium until fully replete would probably need to be conducted in several sessions to study the choices of the rats at different levels of internal sodium to avoid potential confound due to gastric distension. This way, within a session, if rats stop drinking the solutions, we may assume it is because their internal need is satisfied and not because of sickness.

Therefore, we chose to simulate such experimental conditions, by representing the extended two-bottle intake test as a series of short sessions. This different representation also allows us to zoom into the first and last minutes of the long experiments and compare deplete and replete behaviors. **Figure 5** shows example data for the first and the last sessions of each version of the experiment (with NaCl and KCl or with NaCl and LiCl). In Panels A, B and C, the data is collected from a sodium-deplete rat with access to NaCl and KCl, while in Panels D, E and F, the data is collected from a sodium-deplete rat with access to NaCl and LiCl. As we can see in **Figure 5A** during the first session the probability of drinking NaCl increases significantly with sodium deprivation (left). During the last session, the rats are replete, so the choice not to drink anything is the most frequent one (right). In **Figure 5B**, the choices of the rats are represented and in **Figure 5C**, we can see the reward prediction error signals evoked by the outcomes of these choices. We can observe that during the first session, KCl evokes a negative reward prediction error signal because it has an energy cost in our model. This means no DA is released in the NAc, which is consistent with Fortin and Roitman’s result (2018) that KCl does not evoke a DA response in the NAc. During the last session, sodium chloride triggers a weak negative reward prediction error as the rat is replete in sodium. This corresponds to the absence of DA release in the NAc observed by Fortin and Roitman in sodium-replete rats receiving intraoral NaCl infusions (2018). Interestingly, when we simulate an experiment where NaCl and LiCl are available, the probabilities of drinking of NaCl and LiCl remain similar throughout the first and last sessions (**Figure 5D**). We can see the actual simulated choices in **Figure 5E**. We then can also track the reward prediction error signals evoked by the outcomes of these choices (see **Figure 5F**). We note that during the first session, LiCl and NaCl both evoke positive reward prediction error signals, consistent with Fortin and Roitman’s results (2018): intraoral infusions of NaCl and LiCl both trigger DA signals in the NAcs of sodium-deplete rats. Drinking LiCl makes the predicted nutritive value associated with the taste of sodium (and lithium) decrease, which is why after several successive licks of LiCl, the reward prediction error is negative in response to LiCl and NaCl. Drinking NaCl increases the predicted value of the sodium taste, making the taste of sodium more rewarding. Licks of NaCl thus increase the reward prediction error signal. During the last session, the rats have reached the setpoint, so they drink sodium much less often. However, since rats lose some sodium after every trial, their internal sodium level can fall below the setpoint. In that case, NaCl and LiCl evoke a positive reward prediction error signal.

**Figure 5.**
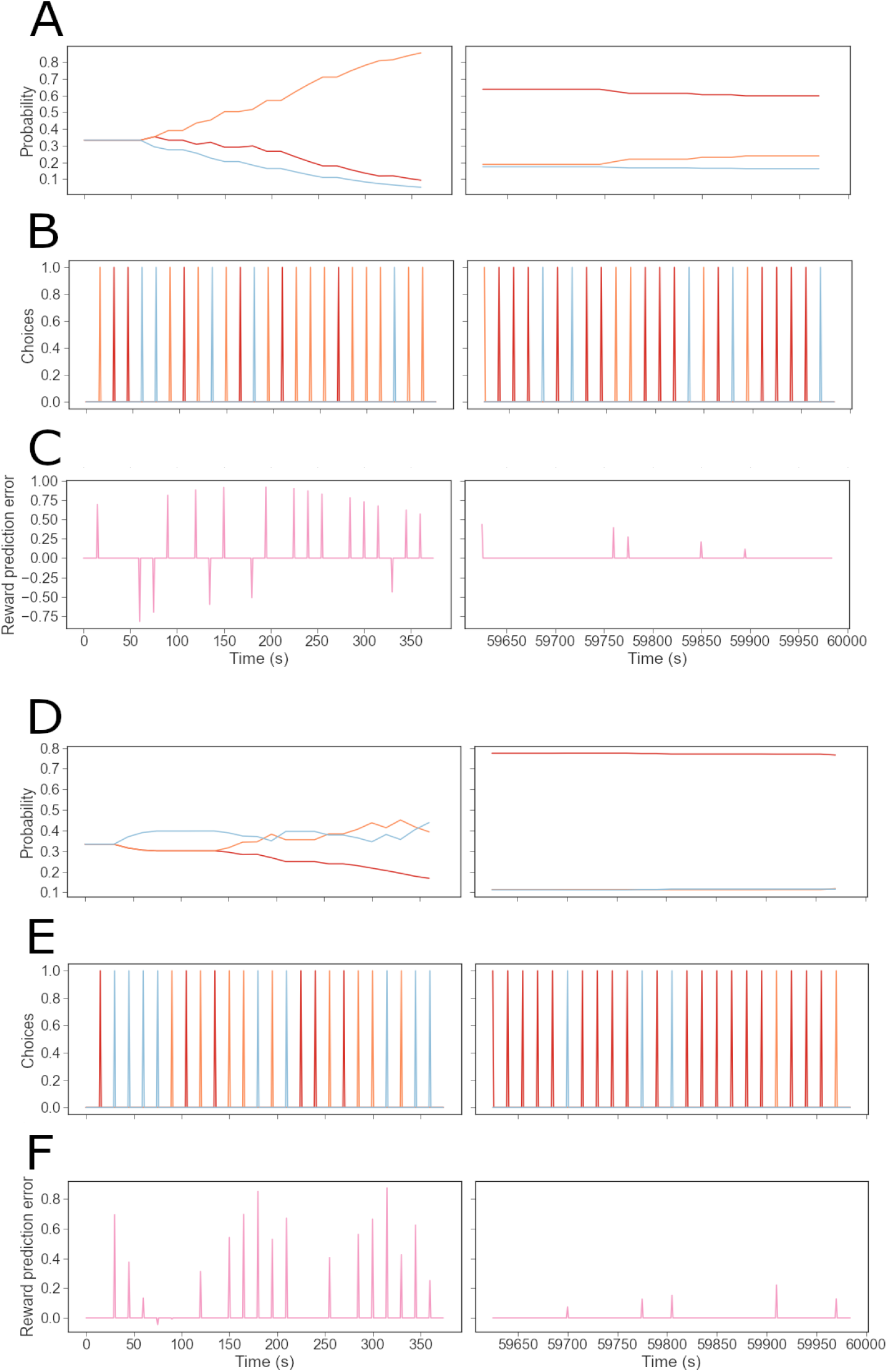
Example data for two simulated rats taking the two-bottle intake test developed by Fortin and Roitman (2018) during an extended period of time, divided into several sessions. Both versions of the experiment last 1000 minutes (4000 trials), divided into 200 sessions of 5 minutes (20 trials). **A,** Evolution of the probability for an individual simulated rat with optimized free parameters to drink NaCl, KCl or nothing over time during the first (left) or last session (right). **B,** choices between drinking NaCl, KCl and nothing over time during the first (left) or last session (right). **C,** Reward prediction error signal over time during the first (left) or last session (right). **D,** Evolution of the probability to drink NaCl, LiCl or nothing over time during the first (left) or last session (right). **E,** choices between drinking NaCl, LiCl and nothing over time during the first (left) or last session (right). **F,** Reward prediction error signal over time during the first (left) or last session (right).

Finally, we wondered what our model would predict for the drinking behavior of rats in presence of only KCl or LiCl. In particular, would sodium-deplete rats keep drinking LiCl, which appears to have the same taste as sodium, to try to satisfy their need? Conversely, would these rats eventually learn that LiCl has no impact on their sodium balance? We used the model to simulate the condition where only KCl or LiCl is available to sodium-deplete rats. Before this experiment, these 2 rats were trained while sodium-deplete as in the experiments described above: the rat with access to KCl could choose from NaCl and KCl and learned not to drink either as it became replete in sodium. A rat with access to LiCl could choose between NaCl and LiCl and also learned not to drink either once it became replete in sodium.

In oder to see clearer what happens in the agent behavior and the corresponding agent internal variable (i.e. the RPE) we show examples of individual behavior and DA dynamics in **Figure 6 F and G**. **Panels 6F** describe the experiment in which only KCl is available. The individual choices are shown in the middle panels. The reward prediction error dynamics, represented in **Figure 6 bottom panels**. As we can see in the top panel, the individual agent’s probability to consume KCl smoothly decreases (null behavior probability decreases) because of the energy cost it imposes without any internal state benefit. Due to this cost, the KCl choice leads to negative RPE signals, which would correspond experimentally to an absence of DA release in the NAc. By the last session, the rat is much more likely to drink nothing (with a probability of about 0.8) than to drink KCl (with a probability of about 0.2). The probability of engaging with KCl is not null even after extended exposure because it represents the probability of making an exploratory decision. During the last session, sodium chloride triggers a weak negative reward prediction error in the model as the agent is replete in sodium. This may correspond to the absence of DA release in the NAc observed by Fortin and Roitman in sodium-replete rats (2018). Panels Figure 6D,E and F describe the experiment in which only LiCl is available. In **Figure 6G**, on the left, we show the evolution of the probability to drink LiCl or nothing over time during the first session. On the right: the probability during the last session. We can see the evolution of the relevant probabilities. We first notice that the probability of drinking LiCl is higher than that of drinking nothing only during the first session and a part of the second one. At the end of the last session, the difference in probability between drinking LiCl and nothing is similar to the one between drinking KCl and nothing, during the last session. Middle panels show the individual choices made. The bottom panels show the reward prediction error dynamics. It appears that, during the first session, the rat learns that LiCl does not increase its internal sodium level and imposes an energy cost. The reward prediction error signals in response to LiCl are first positive and become negative as the rat learns the value of lithium.

**Figure 6.**
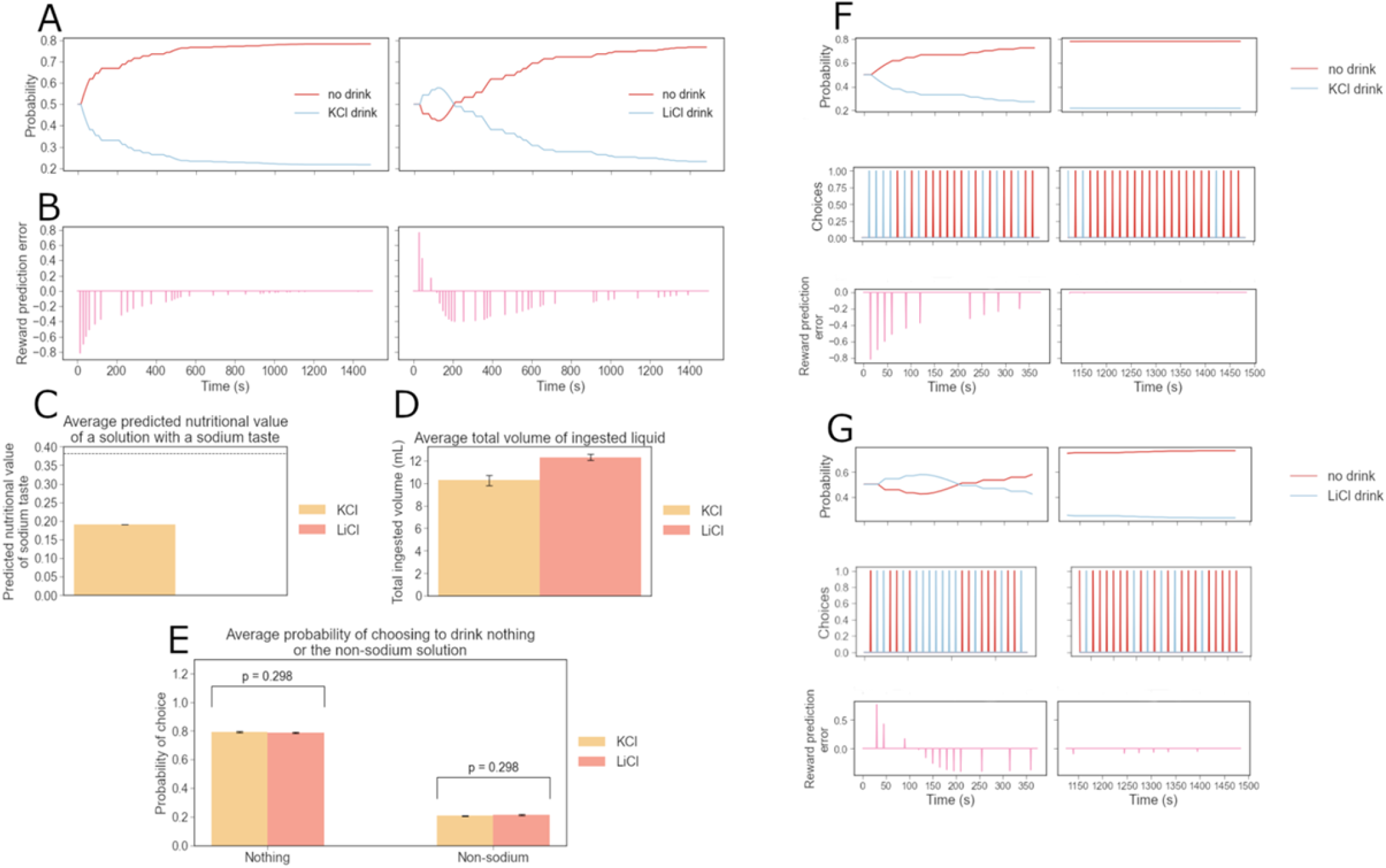
Model predictions on sodium-deplete rat behavior and dopamine dynamics when only non-sodium salt is available. Here we produced example simulated data for an experiment in which two sodium-deplete agents have access to only one salt solution: KCl only and LiCl only. Both versions of the experiment last 25 minutes (100 trials), divided into 5 sessions of 5 minutes (20 trials). A-E give example data for an optimized agent and average data for the experiment in which sodium-depleted agent have access to only one salt solution. For average data, 14 agents have access to KCl while 15 agents have access to LiCl. The free parameter values for the agents in this group (N = 28) are all different and differ slightly from the optimized ones used for the example data. Both versions of the experiment last 25 minutes (100 trials), divided into 5 sessions of 5 minutes (20 trials). We show data for the entire 25-minute experiment. **A,** Evolution of the probability for an agent to drink KCl or nothing over time (left).; probability to drink LiCl (right). **B,** Reward prediction error signals for KCl (left) and LiCl (right). **C,** Average predicted value of a lick of KCl (yellow) or LiCl (orange) based on the taste at the end of the 25-minute experiment. The true nutritive value of one lick of sodium is indicated with a dotted line (p = 2.02e-30). **D**, Average volumes of ingested liquid during the 25-minute simulated experiments: KCl (N = 14) is shown in yellow and LiCl (N = 15) is shown in red (p = 0.001). **E,** Average probabilities to drink the available non-sodium solution (KCl in yellow or LiCl in orange). **F top panels:** Example evolution of the probability of an agent to drink KCl or nothing over time during the first session (left) and the last session (right). **F middle panels:** choices between drinking KCl and nothing over time during the first (left) or last session (right). **F bottom panels:** Reward prediction error signal over time during the first (left) or last session (right). **G top:** Evolution of the probability for a simulated rat with optimized free parameters to drink LiCl or nothing over time during the first (left) or the last session (right). **G middle:** choices between drinking LiCl and nothing over time during the first (left) or last session (right). **G bottom:** Reward prediction error signal over time during the first (left) or last session (right).

**Figure 6** gives example data, represented over the entire duration of the experiments to show evolutions more clearly. It also gives average data that can be tested with future experiments. **Figure 6A** shows the evolution of the probability for the rat to drink KCl or nothing over time (left), and the evolution of the probability for the rat to drink LiCl or nothing over time (right). **Figure 6B** represents the reward prediction error signal of the same rats over time. This different representation gives us a global view of the evolution of the behavior of these individual rats throughout the whole experiment. **Figure 6C** shows the average predicted value of a lick of sodium or lithium based on its taste for rats at the end of the 25-minute experiment. The true nutritive value of one lick of sodium is indicated with a dotted line. This estimate is equal to half of the true value when KCl is available because it does not change in the absence of exposure to NaCl or LiCl. When LiCl is available, this value is close to zero (p = 2.02e-30). **Figure 6E** confirms that at the end of the experiment, rats are on average just as unlikely to drink LiCl (N = 15) as they are to drink KCl (N = 14), which does not taste like NaCl.

This average data allows us to propose an explanation for the individual behavior in **Figure 6A** and **6B**. The agent in Panel A has already learned it is not worth to drink KCl (the action value is initialized with the value the rat previously learned to satisfy its need in sodium). The probability of consuming potassium thus quickly drops to about 0.2. The agent in **Figure 6B** has learned not to drink LiCl as it has kept its associated action value learned while sodium-replete. This agent discovers through an exploratory decision that lithium is rewarding because of the previously learned 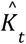, value, but lithium does not increase the internal state, so 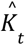 decreases to 0. The reward prediction error signal therefore becomes negative in our model when the agent consumes LiCl, which has a cost but no homeostatic reward. Thus, the probability of consuming lithium decreases quickly decreases to about 0.2. In **Figure 6D**, we chose to represent the total liquid consumed during each version of the experiment. It appears that rats drink more liquid when they have access to LiCl than when KCl is the available solution (p = 0.001).

## Discussion

Here, our goal was to show how sodium directed behaviors and DA responses evoked by sodium and non-sodium stimuli following induction of sodium appetite can be accurately described using a homeostatic reinforcement theoretical framework. To do so, we optimized a simple HRRL agent to learn about the consequences of consuming different sodium (NaCl) and non-sodium (KCl, LiCl)-containing solutions across various states of sodium balance. We show that the behavior of the HRRL model agents closely reproduced experimental data in which sodium depleted rats were given access to NaCl, KCl or LiCl. Much like the rats, following induction of sodium appetite, our HRRL agents strongly preferred solutions that contained sodium or lithium over potassium. We then examined reward prediction error signals on single trials for the HRRL agents in response to consumption of salt-containing solutions. In our HRRL model, the RPE signal should correspond with phasic DA release in the NAc during sodium appetite. We show that the HRRL RPE is strongly modulated by internal state and the properties of the salt stimulus which aligns well with experimentally observed DA signals in the NAc of sodium-deplete rats that were exposed to NaCl, LiCl, and KCl. As in rats with sodium appetite, the HRRL RPE signals were positive for sodium tasting salts (NaCl and LiCl) under deplete conditions and attenuates as the agent approaches a sodium replete state. Furthermore, the HRRL RPE for KCl is negligible which also matches the experimental data.

We then used the HRRL framework to model rat sodium-seeking behavior in longer salt discrimination experiments, where sodium depleted rats are allowed to consume sodium to satiety. Interestingly, our simulated agents began to reduce their consumption of NaCl before becoming sated, suggesting that their behavior reflects anticipation of a future replete state. After extensive experience with salt solutions while sodium depleted, simulated rats that had access to sodium and potassium chloride can predict the nutritive value of one lick of NaCl based on its taste. However, the estimated value of NaCl consumption was halved when simulated rats had access to both sodium and lithium chloride, which have been shown to taste similarly. We finally investigated rat drinking behavior when only lithium chloride is available in the cage, which tastes like sodium chloride but cannot restore lost sodium. Our simulations show that sodium-deprived rats learn that lithium chloride does not fulfill their sodium deficiency and thus stop consuming it. Our simulation results can be tested with experiments which could give further insight on homeostatic state regulation, goal-directed behavior, reinforcement learning, and how these phenomena depend on mesolimbic DA signaling.

In our HRRL framework, we modeled the state of sodium balance through a drive function that represents the deviation between the agents’ current state and an idealized homeostatic set point. In the model, the source of this sodium drive is ambiguous. However, in biological systems, sodium need is sensed via distributed neural systems. There are aldosterone-sensitive neurons in the nucleus of the solitary tract (NTS) that express 11β-hydroxysteroid dehydrogenase type 2 (HSD2+) that are activated by sodium depletion (Geerling et al. 2006). Activation of NTS HSD2+ neurons can drive sodium consumption in sodium replete mice, whereas inhibition reduces sodium consumption in depleted animals (Jarvie and Palmiter 2017). Importantly, NTS HSD2+ neurons project to other hindbrain areas like the parabrachial complex (PB), and pre-locus coeruleus (pre-LC) which contribute to sodium appetite in distinct ways (Geerling and Loewy 2007, Shin et al. 2011). In addition to the NTS, sodium need is also sensed by neurons in the subfornical organ (SFO), which respond to changes in blood osmolality, among other things (Hiyama et al. 2004), and has been shown to regulate sodium appetite (Matusa et al. 2016). Notably, NTS HSD2+ neurons and SFO neurons that respond to osmolality challenges may influence sodium appetite through synergistic influences on neurons in the Bed Nucleus of the Stria Terminalis (Resch et al. 2017). Thus, in biological systems, state sensing -even for a single nutrient- is subject to complex multi-pathway regulation. This is to say nothing of how multiple competing needs interact to influence behavior. In our HRRL models, the drive function was 1-dimensional, as the current state of sodium balance was the only input under consideration.

Future extensions of the HRRL framework could explore how competing drives interact and influence reward computations when multiple nutrients are available (i.e., hunger vs. thirst, blood osmolality and the drive for water vs. sodium). So far we (Keramati & Gutkin 2014; Keramati & Gutkin 2011) and others (Uchida et al 2022) have considered situations where multiple nutrient sources are independent in their impact on the internal state of the agent and therefore the drive function. Interestingly, we did show that under some conditions multiple sources do interact in their impact on shaping behavior in order to satisfy multiple constraints during behavioral policy formation (ref to Juechems). However, neither we nor others have studied what happens when multiple internal states interact through multidimensional drive function. Competition between needs could warp the drive function used to compute the RPE in ways that could not be captured when considering single nutrients. Such simulations could inform further i*n vivo* experiments to probe whether DA signaling evoked by different nutrients or nutrient-predictive cues is shaped by competing drives and whether or not such competition produces non-linear deviations in RPE computations.

In addition to sensing challenges to sodium balance to tune drive states, another key aspect of sodium appetite is the ability to detect sources of sodium in the environment. In the HRRL model, a “tastant” was evaluated by comparing its estimated reward value with the current state of sodium balance. If the agent was below an ideal set point, the tastant was evaluated as positively reinforcing. Otherwise the tastant was evaluated negatively as it further exacerbated deviations from homeostasis and cost energy to approach and consume. *In vivo*, sensing sodium primarily begins via Na+ selective epithelial sodium channels (eNACs) which are critical for sodium detection and discrimination (Geran and Spector 2000; Smith et al. 2012; but see Treesukosol et al. 2007). Sodium taste information is relayed to the brain via the chorda tympani (CT; O’Keefe, Schumm, and Smith 1994; Golden et al. 2012) and CT transection disrupts expression of sodium appetite (Sollars and Bernstein 1992; Breslin, Spector, and Grill 1993). Interestingly, CT responses to sodium solutions are augmented by sodium appetite (Contreras and Frank, 1979), suggesting that changes in sodium balance can exert widespread effects on taste processing, even outside the CNS. In the CNS, sodium appetite alters sodium taste responses in numerous brain areas, including hindbrain structures like the NTS (Jacobs, Mark, and Scott 1988; McCaughey and Scott 2000) and PB (Shimura, Komori, and Yamamoto 1997) as well as circuits linked to reinforcement learning like the striatum (Loriaux, Roitman, and Roitman 2011) and the mesolimbic DA system (Cone et al. 2016; Fortin and Roitman 2018). Such widespread changes in sodium taste processing likely contribute to the robust changes in consummatory responses and affective orosensory evaluation of sodium containing solutions typically observed in sodium appetite (Breslin et al. 1993; Berridge et al. 1984).

Linking the gustatory properties of sodium-containing stimuli (taste, phasic) with sodium need (state, tonic) is essential for sodium appetite. Changes in physiological state (tonic) serve to bias the organisms behavioral repertoire towards sodium-seeking behaviors such that sodium intake is invigorated when tastants that contain sodium are identified. For example, consummatory responses such as bursts of licking, which are thought to reflect the palatability of a taste stimulus (Johnson et al. 2018), increase following sodium depletion (Breslin et al. 1993). Prior work suggests that taste sensing and state interact in an additive fashion (Breslin et al. 1993), but if either is disrupted, expression of sodium appetite is impaired (Bernstein and Hennessy 1987; Jarvie and Palmiter 2017). Perhaps the key feature of the HRRL framework is that by incorporating a drive function that tracks deviations from homeostasis into a traditional RL-based agent, this enabled us to model the taste-state interactions that drive both current and future sodium-seeking behaviors associated with sodium appetite. At the moment of consumption (e.g., when an RPE is triggered), it is likely that state-dependent changes in taste information is relayed to DA neurons via the PB and preLC, both of which innervate the VTA (Shin et al. 2011). PB neurons express cFos following sodium consumption in sodium-deplete rats and this corresponds with reduced cFos staining in NTS HSD2+ neurons (Geerling and Loewy 2007). Moreover, preLC FoxP2+ neurons that receive efferents from the NTS are strongly implicated in sodium appetite (Lee et al. 2019) and project to the VTA (Fortin and Roitman 2018). By tuning RPEs according to the current state of the animal, the HRRL model may mimic PB and preLC influences on VTA DA signaling that track physiological need.

Future work could explore whether the HRRL model also captures longer term plasticity in reinforcement circuits associated with sodium appetite. For example, sodium appetite is augmented by a single prior experience with sodium depletion (Sakai et al. 1987), while repeated depletions increase need free sodium intake long after sodium levels are replenished (Sakai et al. 1989; Deitz, Curtis, and Contreras 2005). In terms of RPE signaling, mesolimbic DA neurons appear to encode sodium cues as reward predictive only after extensive experience with cue-sodium pairings while sodium deplete (Cone et al. 2016). However, after such associations are learned, phasic DA release to the sodium cue is only observed when the animals are in a state of need (Cone et al. 2016). It is likely the HRRL model would already account for this in its underlying math. Another challenging issue is how the proposed drive function is encoded in neural activity and how multiple drives may be prioritized with respect to each other (see also the discussion on multiple constraint satisfaction above). Juechems and colleagues (Juechems et al., 2018) have suggested that the rostral anterior cingulate cortex (rACC) encodes the imbalance between assets such as food, water or money. This information can be used to decide which internal need should be satisfied first. The ventromedial prefrontal cortex (vmPFC) encodes how much a given choice brings the internal needs back to a balanced state. These regions thus link information on the internal states with the valuation of actions and couple be the potential neural systems for drive representation.

An interesting point to consider in light of our results is whether the taste can be considered as another CS whose predicted impact of the consumed US on the internal state is learned. A study on DA in mice consuming sucrose draws conclusions supporting this hypothesis (Beeler et al., 2012). According to this study, a taste evokes a DA response only if the animal has associated through prior learning this taste with a nutritional value. In other words, the hedonic property of taste cannot alone be responsible for reinforcement. The study suggests that taste acts as a conditioned stimulus predictive of a food reward. This implies that the association between a taste and a nutritional value can be extinguished by giving the animal extensive experience with the taste in the absence of a value (an artificial sweetener instead of sucrose, for example). When it comes to sodium, however, taste does not seem to act simply as a conditioned stimulus and appetite appears to be regulated by a more complex interplay of both innate and learned mechanisms. This is reflected in our model by the non-linear drive function and the non-zero initial value of 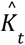. However, in our simulations, it is possible to extinguish seeking behavior towards the sodium taste after exposure of sodium-deplete rats to LiCl. This suggests that the subjective value attached to sodium taste can be modified with learning, which should be investigated with new experiments.

Intriguingly a recent proposal suggests that gustatory information could act as a hedonic predictor of the long-run worth of a good (Dayan 2022). This suggestion appears at a first glance compatible with our work, for example just as with the gustatory hedonic” account in our simulation the gustatory “reward/value” rapidly leads to motivate consumption behavior and is ultimately dominated by the nutritive lack of impact for LiCl with the end value converging to zero.

Our model, as all computational models, faces several limitations: the actions are necessarily chosen and performed by the agent at discrete and regular time steps. Moreover, the solutions are consumed at a fixed volume. A continuous time model, in which the internal state of the agent has to be constantly regulated with actions that maintain its homeostasis, would be closer to reality. Indeed, the temporality of physiological regulation is important as the deviation from the setpoint must be reduced as soon as possible. We have previously shown that taking the shortest path to reach the setpoint optimizes fitness and that HRRL agents learn to preemptively avoid life-threatening excursions far from the homeostatic optima (Keramati and Gutkin, 2014). The proposed framework remains quite an abstract algorithmic model: how it relates to the neural circuitry mechanistically remains to be studied. For example, as discussed above the circuits that mediate taste-state interactions in the brain are multi-faceted and could be communicated with the DA system through many convergent pathways. Another challenge that remains is how might one observe the drive function? A hint of predictive internal state encoding has been shown experimentally (see discussion above and Livneh et al. 2017), however it is not clear if such neural traces represent the internal state itself or the drive function necessary for learning and generating the behaviors. In fact, perhaps the drive function properties could be a key to individual differences in homeostatically motivated learned behaviors. Understanding how one may measure the properties of the drive function from behavioral observations thus remains a key challenge.

In conclusion, we have shown that a homeostatic reinforcement learning theory can account for behaviors motivated by sodium appetite. A recent study also put forth a similar argument (REF) - focusing on a different dataset, they showed that HRRL agents can match two-bottle test behavior for sodium and water choices. Here, we extend these ideas to show that HRRL can also account quantitatively for the taste- and state-dependent DA responses, arguing that such DA signaling may in fact be causal for the learning of such appetitive behaviors.

## Acknowledgments

This work was supported by NIH grant DA0256234 (MFR); ANR grants ANR-17-EURE-0017 and ANR-10-IDEX-0001-02 (BSG).

## References

1. Tian J, Huang R, Cohen JY, Osakada F, Kobak D, Machens CK, Callaway EM, Uchida N, Watabe-Uchida M. 2016 Distributed and Mixed Information in Monosynaptic Inputs to Dopamine Neurons. Neuron. 91(6):1374–1389. doi: 10.1016/j.neuron.2016.08.018. Epub 2016 Sep 8. PMID: 27618675; PMCID: PMC5033723.

2. Schultz W. Multiple reward signals in the brain. 2000 Nat Rev Neurosci. 1(3):199–207. doi: 10.1038/35044563. PMID: 11257908.

3. Livneh, Y., Ramesh, R., Burgess, C. et al. 2017 Homeostatic circuits selectively gate food cue responses in insular cortex. Nature 546:611–616. https://doi.org/10.1038/nature22375

4. Geerling JC, Loewy AD. 2008 Central regulation of sodium appetite. Exp Physiol. 93(2):177–209. doi: 10.1113/expphysiol.2007.039891. Epub 2007 Nov 2. PMID: 17981930.

5. Augustine V, Lee S, Oka Y. 2020 Neural Control and Modulation of Thirst, Sodium Appetite, and Hunger. Cell. 180(1):25–32. doi: 10.1016/j.cell.2019.11.040. PMID: 31923398; PMCID: PMC7406138.

6. Schultz W 1998 Predictive Reward Signal of Dopamine Neurons Journal of Neurophysiology 80(1): 1–27

7. Keiflin R, Janak PH. 2015 Dopamine Prediction Errors in Reward Learning and Addiction: From Theory to Neural Circuitry. Neuron. 88(2):247–63. doi: 10.1016/j.neuron.2015.08.037.

8. Dabney, W, Kurth-Nelson, Z., Uchida, N. et al. 2020 A distributional code for value in dopamine-based reinforcement learning. Nature 577:671–675. https://doi.org/10.1038/s41586-019-1924-6

9. Kutlu MG, Zachry JE, Melugin PR, Cajigas SA, Chevee MF, Kelly SJ, Kutlu B, Tian L, Siciliano CA, Calipari ES. 2021 Dopamine release in the nucleus accumbens core signals perceived saliency. Curr Biol. 31(21):4748–4761.e8. doi: 10.1016/j.cub.2021.08.052. Epub 2021 Sep

10. Liu S, Borgland SL. 2015 Regulation of the mesolimbic dopamine circuit by feeding peptides. Neuroscience. 289:19–42. doi: 10.1016/j.neuroscience.2014.12.046

11. Cone JJ, Roitman JD, Roitman MF. 2015 Ghrelin regulates phasic dopamine and nucleus accumbens signaling evoked by food-predictive stimuli. J Neurochem. 133(6):844–56. doi: 10.1111/jnc.13080.

12. Mietlicki-Baase EG, Reiner DJ, Cone JJ, Olivos DR, McGrath LE, Zimmer DJ, Roitman MF, Hayes MR. 2015 Amylin modulates the mesolimbic dopamine system to control energy balance. Neuropsychopharmacology. 40(2):372–85. doi: 10.1038/npp.2014.180.

13. Mebel DM, Wong JC, Dong YJ, Borgland SL. 2012 Insulin in the ventral tegmental area reduces hedonic feeding and suppresses dopamine concentration via increased reuptake. Eur J Neurosci. 36(3):2336–46. doi: 10.1111/j.1460-9568.2012.08168.x.

14. Cone JJ, McCutcheon JE, Roitman MF. 2014 Ghrelin acts as an interface between physiological state and phasic dopamine signaling. J Neurosci. 34(14):4905–13. doi: 10.1523/JNEUROSCI.4404-13.2014

15. Cone JJ, Fortin SM, McHenry JA, Stuber GD, McCutcheon JE, Roitman MF. 2016 Physiological state gates acquisition and expression of mesolimbic reward prediction signals. Proc Natl Acad Sci U S A. 113(7):1943–8. doi: 10.1073/pnas.1519643113

16. Fortin SM, Roitman MF. 2018 Challenges to Body Fluid Homeostasis Differentially Recruit Phasic Dopamine Signaling in a Taste-Selective Manner. J Neurosci. 38(31):6841–6853. doi: 10.1523/JNEUROSCI.0399-18.2018

17. Hsu, T. M., Bazzino, P., Hurh, S. J., Konanur, V. R., Roitman, J. D., & Roitman, M. F. 2020 Thirst recruits phasic dopamine signaling through subfornical organ neurons. Proceedings of the National Academy of Sciences of the United States of America, 117(48):30744–30754. https://doi.org/10.1073/pnas.2009233117

18. Grove, J.C.R., Gray, L.A., La Santa Medina, N. et al. 2022 Dopamine subsystems that track internal states. Nature 608:374–380, https://doi.org/10.1038/s41586-022-04954-0

19. Reichenbach, A., Clarke, R. E., Stark, R., Lockie, S. H., Mequinion, M., Dempsey, H., Rawlinson, S., Reed, F., Sepehrizadeh, T., DeVeer, M., Munder, A. C., Nunez-Iglesias, J., Spanswick, D. C., Mynatt, R., Kravitz, A. V., Dayas, C. V., Brown, R., & Andrews, Z. B. 2022 Metabolic sensing in AgRP neurons integrates homeostatic state with dopamine signalling in the striatum. eLife, 11, e72668. https://doi.org/10.7554/eLife.72668

20. Denton D 1982 The hunger for salt: an anthropological, physiological, and medical analysis Berlin; New York: Springer-Verlag, 1982. ISBN 0387112863

21. Curt P. Richter 1936 INCREASED SALT APPETITE IN ADRENALECTOMIZED RATS American Journal of Physiology-Legacy Content 115:1, 155–161

22. Nachman, M. (1963). Learned aversion to the taste of lithium chloride and generalization to other salts. Journal of Comparative and Physiological Psychology, 56(2), 343–349. https://doi.org/10.1037/h0046484

23. Contreras RJ, Frank M. 1979 Sodium deprivation alters neural responses to gustatory stimuli. J Gen Physiol. 73(5):569–94. doi: 10.1085/jgp.73.5.569.

24. Hull CL 1943 Principles of behavior: an introduction to behavior theory New York: Appleton-Century-Crofts. Mehdi Keramati Boris Gutkin 2014 Homeostatic reinforcement learning for integrating reward collection and physiological stability eLife 3:e04811.

25. Keramati, M., Durand, A., Girardeau, P., Gutkin, B., & Ahmed, S. H. 2017 Cocaine addiction as a homeostatic reinforcement learning disorder. Psychological Review, 124(2):130–153. https://doi.org/10.1037/rev0000046

26. Gonçalves PJ, Lueckmann JM, Deistler M, Nonnenmacher M, Öcal K, Bassetto G et al. 2020 Training deep neural density estimators to identify mechanistic models of neural dynamics, eLife 9:e56261

27. Geerling JC, Engeland WC, Kawata M, Loewy AD. 2006 Aldosterone target neurons in the nucleus tractus solitarius drive sodium appetite. J Neurosci. 26(2):411–7. doi: 10.1523/JNEUROSCI.3115-05.2006

28. Jarvie, B., Palmiter, R. 2017 HSD2 neurons in the hindbrain drive sodium appetite. Nature Neuroscience 20:167–169 https://doi.org/10.1038/nn.4451

29. Geerling JC, Loewy AD. 2007 Sodium deprivation and salt intake activate separate neuronal subpopulations in the nucleus of the solitary tract and the parabrachial complex. J Comp Neurol. 504(4):379–403. doi: 10.1002/cne.21452

30. Shin JW, Geerling JC, Stein MK, Miller RL, Loewy AD. 2011 FoxP2 brainstem neurons project to sodium appetite regulatory sites. J Chem Neuroanat. 42(1):1–23. doi: 10.1016/j.jchemneu.2011.05.003.

31. Hiyama TY, Watanabe E, Okado H, Noda M. 2004 The subfornical organ is the primary locus of sodium-level sensing by Na(x) sodium channels for the control of salt-intake behavior. J Neurosci. 24(42):9276–81. doi: 10.1523/JNEUROSCI.2795-04.2004.

32. Matsuda, T., Hiyama, T., Niimura, F. et al. 2017 Distinct neural mechanisms for the control of thirst and salt appetite in the subfornical organ. Nat Neurosci 20:230–241 https://doi.org/10.1038/nn.4463

33. Resch JM, Fenselau H, Madara JC, Wu C, Campbell JN, Lyubetskaya A, Dawes BA, Tsai LT, Li MM, Livneh Y, Ke Q, Kang PM, Fejes-Tóth G, Náray-Fejes-Tóth A, Geerling JC, Lowell BB. 2017 Aldosterone-Sensing Neurons in the NTS Exhibit State-Dependent Pacemaker Activity and Drive Sodium Appetite via Synergy with Angiotensin II Signaling. Neuron. 96(1):190–206.e7. doi: 10.1016/j.neuron.2017.09.014

34. Uchida Y, Hikida T, Yamashita Y. 2022 Computational Mechanisms of Osmoregulation: A Reinforcement Learning Model for Sodium Appetite. Front Neurosci. 16:857009. doi: 10.3389/fnins.2022.857009.

35. Keramati, M and Gutkin, B.S. 2011 A Reinforcement Learning Theory for Homeostatic Regulation. Advances in Neural Information Systems 24 J. Shawe-Taylor and R. Zemel and P. Bartlett and F. Pereira and K.Q. Weinberger (eds) ISBN: 9781618395993

36. Geran, L. C., & Spector, A. C. 2000 Sodium taste detectability in rats is dependent of anion size: The psychophysical characteristics of the transcellular sodium taste transduction pathway. Behavioral Neuroscience, 114(6):1229–1238. https://doi.org/10.1037/0735-7044.114.6.1229

37. Kimberly R. Smith, Yada Treesukosol, A. Brennan Paedae, Robert J. Contreras, and Alan C. Spector 2012 Contribution of the TRPV1 channel to salt taste quality in mice as assessed by conditioned taste aversion generalization and chorda tympani nerve responses American Journal of Physiology-Regulatory, Integrative and Comparative Physiology 303:11, R1195–R1205

38. Yada Treesukosol, Vijay Lyall, Gerard L. Heck, John A. DeSimone, and Alan C. Spector 2007 A psychophysical and electrophysiological analysis of salt taste in *Trpv1* null mice American Journal of Physiology-Regulatory, Integrative and Comparative Physiology 292:5, R1799–R1809

39. O’Keefe GB, Schumm J, Smith JC. 1994 Loss of sensitivity to low concentrations of NaCl following bilateral chorda tympani nerve sections in rats. Chem Senses. 19(2):169–84. doi: 10.1093/chemse/19.2.169.

40. Golden GJ, Ishiwatari Y, Theodorides ML, Bachmanov AA. 2011 Effect of chorda tympani nerve transection on salt taste perception in mice. Chem Senses. 36(9):811–9. doi: 10.1093/chemse/bjr056.

41. Sollars, S. I., & Bernstein, I. L. 1992 Sodium appetite after transection of the chorda tympani nerve in Wistar and Fischer 344 rats. Behavioral Neuroscience, 106(6):1023–1027. https://doi.org/10.1037/0735-7044.106.6.1023

42. P. A. Breslin, A. C. Spector, and H. J. Grill 1993 Chorda tympani section decreases the cation specificity of depletion-induced sodium appetite in rats American Journal of Physiology-Regulatory, Integrative and Comparative Physiology 264:2, R319–R323

43. Contreras RJ, Frank M. 1979 Sodium deprivation alters neural responses to gustatory stimuli. J Gen Physiol. 73(5):569–94. doi: 10.1085/jgp.73.5.569.

44. Jacobs KM, Mark GP, Scott TR. 1988 Taste responses in the nucleus tractus solitarius of sodium-deprived rats. J Physiol. 406:393–410. doi: 10.1113/jphysiol.1988.sp017387.

45. Stuart A. McCaughey and Thomas R. Scott 2000 Rapid induction of sodium appetite modifies taste-evoked activity in the rat nucleus of the solitary tract American Journal of Physiology-Regulatory, Integrative and Comparative Physiology 279:3, R1121–R1131

46. Shimura T, Komori M, Yamamoto T. 1997 Acute sodium deficiency reduces gustatory responsiveness to NaCl in the parabrachial nucleus of rats. Neurosci Lett. 236(1):33–6. doi: 10.1016/s0304-3940(97)00745-3

47. Loriaux AL, Roitman JD, Roitman MF. 2011 Nucleus accumbens shell, but not core, tracks motivational value of salt. J Neurophysiol. 106(3):1537–44. doi: 10.1152/jn.00153.2011.

48. P. A. Breslin, J. M. Kaplan, A. C. Spector, C. M. Zambito, and H. J. Grill 1993 Lick rate analysis of sodium taste-state combinations American Journal of Physiology-Regulatory, Integrative and Comparative Physiology 264:2, R312–R318

49. Berridge, K. C., Flynn, F. W., Schulkin, J., & Grill, H. J. 1984 Sodium depletion enhances salt palatability in rats. Behavioral Neuroscience, 98(4):652–660, https://doi.org/10.1037/0735-7044.98.4.652

50. Johnson, A.W. 2018 Characterizing ingestive behavior through licking microstructure: Underlying neurobiology and its use in the study of obesity in animal models. International Journal of Developmental Neuroscience, 64: 38–47. https://doi.org/10.1016/j.ijdevneu.2017.06.012

51. Bernstein IL, Hennessy CJ. 1987 Amiloride-sensitive sodium channels and expression of sodium appetite in rats. Am J Physiol. 253(2 Pt 2):R371–4. doi: 10.1152/ajpregu.1987.253.2.R371.

52. Jarvie, B., Palmiter, R. 2017 HSD2 neurons in the hindbrain drive sodium appetite. Nat Neurosci 20:167–169, https://doi.org/10.1038/nn.44517

53. Shin JW, Geerling JC, Stein MK, Miller RL, Loewy AD. 2011 FoxP2 brainstem neurons project to sodium appetite regulatory sites. J Chem Neuroanat. 42(1):1–23. doi: 10.1016/j.jchemneu.2011.05.003.

54. Lee, S., Augustine, V., Zhao, Y. et al. 2019 Chemosensory modulation of neural circuits for sodium appetite. Nature 568:93–97, https://doi.org/10.1038/s41586-019-1053-2

55. Sakai RR, Fine WB, Epstein AN, Frankmann SP. 1987 Salt appetite is enhanced by one prior episode of sodium depletion in the rat. Behav Neurosci. 101(5):724–31. doi: 10.1037//0735-7044.101.5.724.

56. Sakai, R. R., Frankmann, S. P., Fine, W. B., & Epstein, A. N. 1989 Prior episodes of sodium depletion increase the need-free sodium intake of the rat. Behavioral Neuroscience, 103(1):186–192. https://doi.org/10.1037/0735-7044.103.1.186

57. Dietz DM, Curtis KS, Contreras RJ 2006 Taste, Salience, and Increased NaCl Ingestion after Repeated Sodium Depletions, Chemical Senses, 31(1):33–41, https://doi.org/10.1093/chemse/bjj003

58. Juechems K, Summerfield C. Where Does Value Come From? Trends Cogn Sci. 2019 Oct;23(10):836–850. doi: 10.1016/j.tics.2019.07.012

59. Dayan P (2022) “Liking” as an early and editable draft of long-run affective value. PLoS Biol 20(1): e3001476. https://doi.org/10.1371/journal.pbio.3001476

